# Dynamic Inter-Modality Source Coupling Reveals Sex Differences in Children based on Brain Structural-Functional Network Connectivity: A Multimodal MRI Study of the ABCD Dataset

**DOI:** 10.1101/2025.07.23.666366

**Authors:** A. Kotoski, S-L. Wiafe, J. M. Stephen, Y-P. Wang, T. W. Wilson, V. D. Calhoun

## Abstract

**Background:** Sex differences in brain development are well-documented, yet the dynamic coupling between structure and function remains underexplored. We introduce dynamic inter-modality source coupling (dIMSC), extending our previous work to link structural MRI source-based morphometry (SBM) with dynamic functional network connectivity (dFNC).

**Methods:** We used data from 500 participants in the Adolescent Brain Cognitive Development (ABCD) study (ages 9-11) and combined SBM-derived gray matter sources with sliding-window dFNC. dIMSC was computed as the time-resolved cross-correlation between these modalities to quantify structure-function coupling strength. We evaluated sex differences in these coupling profiles and used a cognitive moderation analysis to validate the sensitivity of the identified coupling states.

**Results:** Significant sex-specific patterns emerged: males exhibited stronger positive coupling in sensorimotor regions (postcentral gyrus), while females showed stronger coupling in higher-order associative regions (inferior parietal lobule). These configurations were functionally distinct: higher positive coupling predicted better crystallized cognition (vocabulary) in females, whereas it predicted better fluid cognition (working memory) in males.

**Conclusion:** These findings validate the functional sensitivity of dIMSC, demonstrating that the time-resolved coupling states it identifies are neurobiologically meaningful and differentially predictive of cognitive performance as a function of biological sex, consistent with known sex differences in neurodevelopmental during late childhood.

**Author Summary:** The brain is not static, its activity constantly shifts, and how closely that activity follows the brain’s physical structure changes moment to moment. Most studies examine structure-function relationships as fixed averages, potentially missing important temporal dynamics. Here, we applied a novel dynamic approach to characterize how brain structure and function couple and decouple over time in 500 children from the ABCD Study, and whether this varies by sex. We found that males and females differ meaningfully in how much time specific brain networks spend in different coupling states. Crucially, these differences carry cognitive consequences: in females, stronger structural anchoring of higher-order associative regions predicted better crystallized cognition, while in males it predicted better fluid reasoning.

## I. INTRODUCTION

Sex differences in brain development, historically underexamined in neuroscience, are now receiving increased attention due to their potential role in shaping cognitive, behavioral, and psychiatric trajectories (Bangasser & Valentino, 2014; Miller & Halpern, 2014; Okami & Shackelford, 2001). From early childhood through adolescence, males and females exhibit differences in brain maturation rates, cortical thickness, connectivity, and functional organization (De Bellis et al., 2001; Gong et al., 2011). Prior studies have reported sex-specific patterns in brain volume, white matter development, and resting-state network activity (Alarcón et al., 2015; De Bellis et al., 2001; Perrin et al., 2009), with some suggesting that these differences may influence emotional regulation, or susceptibility to disorders such as attention-deficit/hyperactivity disorder, anxiety, and autism (Arnold, 1996; Baron-Cohen et al., 2005; Gardener et al., 2013). Importantly, these reported differences reflect group-level means, and within-sex variability is often substantial, in some cases exceeding the magnitude of between-sex differences (Bottenhorn et al., 2023; Torgerson et al., 2025; Wierenga et al., 2018). However, while many of these findings highlight brain differences between sexes, most of the literature has examined brain structure and function in isolation, with limited insight into how brain structure-function interact across sexes during development. Critically, these sex differences in brain organization are not merely anatomical, they carry functional consequences for cognition. Males and females show well-documented differences in cognitive performance, including female advantages in verbal and crystallized cognition and male advantages in visuospatial and fluid reasoning (Miller & Halpern, 2014), yet the neural mechanisms linking sex-specific brain organization to these cognitive outcomes remain poorly understood. Therefore, the mechanisms by which structure-function coupling differs between males and females, and how those differences relate to cognitive development, remain largely unexplored.

Structural MRI (sMRI) and resting-state functional MRI (rs-fMRI) are widely used to study the brain during development and to evaluate sex differences (De Bellis et al., 2001; Gong et al., 2011). sMRI provides information into gray and white matter anatomy, including cortical volume, thickness, and surface area (Phan et al., 2021). Source-based morphometry (SBM) is a multivariate extension of sMRI analysis that applies independent component analysis to gray matter volume maps, identifying spatially independent patterns of covarying structural variation across individuals and quantifying their expression in each subject as a loading parameter (Gupta et al., 2019). Functional MRI (fMRI) is used to indirectly study the brain’s activity by measuring the changes in blood-oxygen levels (Correa et al., 2009). rs-fMRI applies this technique at rest capturing the connectivity of brain regions when no task or stimulus is being applied (Lee et al., 2013).

Prior research has demonstrated sex-related differences in both sMRI and fMRI, particularly within the developmental window of late childhood and adolescence (Giedd et al., 1999; Giedd et al., 1997; Satterthwaite et al., 2015). Structural studies frequently report that males show larger total brain volumes, after correcting for total intracranial volume, while females exhibit accelerated maturation and achieve peak cortical volume earlier (Ingalhalikar et al., 2014; Lenroot & Giedd, 2010; Shaywitz et al., 1995). Regionally, females often show larger volumes in areas related to language and associative cortex (Gur et al., 1999; Harasty et al., 1997). Resting-state studies have revealed sex differences in functional network connectivity, particularly in the default-mode network, dorsal attention network, and salience network (Shaywitz et al., 1995). dFNC studies have also shown sex-specific temporal characteristics, where females tend to transition more rapidly between network configurations, and males exhibit longer periods of stability within specific states (Murray et al., 2021; Nini et al., 2017). There is also existing work in structure-function coupling suggesting sex differences in how brain structure constrains functional connectivity in adult samples (Wan et al., 2022; Zhao et al., 2021). Importantly, these structural and functional differences align with the cognitive domains in which sex differences are most consistently reported: the regions where females show earlier maturation and greater volume are precisely those that support verbal memory, reading, and crystallized cognition [16, 20, 21]. Conversely, the more prolonged maturation trajectory observed in males may support the development of fluid reasoning and visuospatial abilities [1, 14]. These convergent patterns suggest that sex differences in structure-function coupling may represent a mechanistic bridge between brain organization and cognitive development.

To address the question of how structure-function relationship may differ by sex, it is crucial to investigate their short-term dynamic interplay. Neurobiological processes underlying development, such as synaptic pruning, myelination, and activity-dependent plasticity, are continuous and time-varying, and these processes are known to exhibit sex-specific timing, demonstrated in human neuroimaging studies (De Bellis et al., 2001; Giedd et al., 1999; Lenroot & Giedd, 2006; Sisk & Zehr, 2005). Therefore, a static, time-averaged assessment of structure-function coupling is inherently limited, not because a single scan can capture ongoing maturational processes such as synaptic pruning or myelination, but because averaging across the entire scan obscures the moment-to-moment variability in how functional dynamics align with structural architecture. The neurodevelopmental relevance of dynamic FC lies in its sensitivity to individual differences in brain state organization, children at different stages of maturation, or with different sex-typical developmental trajectories, may differ in how frequently and how strongly their functional dynamics align with structural constraints across the scan session (Chen et al., 2024; Icer et al., 2020). These transient periods of alignment, captured by dIMSC, may therefore serve as a more sensitive index of sex-specific organizational features than static coupling, even though the underlying maturational processes unfold over much longer timescales. Moreover, these transient states of structural alignment are not merely technical artefacts, they may represent the precise moments when cognitive operations are most efficiently supported by the underlying anatomy, and understanding how often a brain dwells in these states may reveal sex-specific neural strategies for cognition. By capturing the temporal dimension of the scan session, a dynamic approach is necessary to reveal the moments when structural constraints align with or oppose functional network activity, offering a more sensitive index of sex differences in ongoing neurodevelopmental trajectories.

In a prior study, we introduced inter-modality source coupling (IMSC) (Kotoski et al., 2024), a method that quantified the association between structural and functional brain components derived from independent component analysis (ICA) (Lee, 1998). While IMSC has provided insights into static structure-function coupling, it does not account for temporal fluctuations that may reflect dynamic brain activity. To address this limitation, we propose a novel extension called dynamic inter-modality source coupling (dIMSC) which allows for time-resolved assessment of structure-function coupling.

The neurobiological hypothesis guiding the use of dIMSC is that functional network dynamics are not uniformly constrained by underlying structural architecture. To capture the temporal evolution of brain function, we use dynamic functional network connectivity (dFNC) derived from sliding-window based Pearson correlation, which estimates time-varying connectivity between functional brain networks (Allen et al., 2014; Wiafe et al., 2024). dFNC provides a more nuanced representation of brain activity by capturing transient brain states that are obscured when averaging over longer periods, as is done in static FNC. Some of these dynamic states may be more closely aligned with underlying brain structure than others, making dFNC a more sensitive measure for detecting structure-function coupling. For brain structure, we utilize the source-based morphometry (SBM) components, spatially independent gray matter sources derived from ICA, to capture patterns of covarying structural variation across individuals (Xu et al., 2009). We selected SBM over other structural modalities because fMRI signals originate within gray matter (Logothetis et al., 2001). Therefore, SBM provides the direct anatomical substrate of the functional networks we are analyzing (Segall et al., 2012), allowing us to test how the structural integrity of these cortical nodes constrains their dynamic functional pattern (Sui et al., 2014). dIMSC is computed by correlating SBM components with each temporal snapshot of dFNC pattern. The present study aims to: (1) characterize sex differences in dynamic structure-function coupling across 53 brain components in children aged 9-11 years, and (2) validate the functional sensitivity of dIMSC by demonstrating that the identified coupling states are differentially predictive of cognitive performance as a function of biological sex, establishing cognitive relevance as a critical proof-of-concept for the method. Based on this literature, we formulate two testable hypotheses. First, we hypothesize that females will show stronger positive (structurally-anchored) coupling in higher-order associative and language-related regions, consistent with their earlier structural maturation and proportionally larger volumes in these areas, whereas males will show stronger positive coupling in sensorimotor and visuospatial regions, consistent with a more prolonged, sensorimotor-dominant structural trajectory. Second, because these coupling states are expected to carry direct cognitive relevance, we hypothesize that dIMSC occupancy in structurally-anchored associative regions will more strongly predict crystallized and verbal cognition in females, whereas dIMSC occupancy in structurally-anchored sensorimotor regions will more strongly predict fluid reasoning and executive function in males. Confirmation of this sex-specific correspondence between coupling location and cognitive domain would provide direct evidence that the coupling states identified by dIMSC are neurobiologically meaningful rather than statistical artifacts.

## II. METHODS

In this study, we use a novel method to examine time-varying interactions between brain structure and function in children aged 9–11 years-old. We analyzed baseline data from the Adolescent Brain Cognitive Development (ABCD) study, which includes over 11,000 children recruited from 21 sites across the United States. We used ABCD Data Release 5.1. Consistent with ABCD Study data-quality guidance regarding a documented intermittent scanner issue at Site 15 affecting functional and diffusion MRI data acquired prior to August 2018, we confirmed during subject selection that no participants included in our final analytic sample of 500 individuals were scanned at Site 15 prior to this date. The sample is demographically diverse and was designed to approximate the U.S. population in terms of age, sex, ethnicity, and socioeconomic status (Volkow et al., 2018). This approach captures transient coupling between dynamic functional network connectivity (dFNC) and structural MRI gray matter volume. We then use it to explore sex differences in dFNC-sMRI coupling across brain regions during this critical developmental period.

### A. Data

To reduce computational demand, we selected a random subset of 500 unrelated individuals from the ABCD baseline sample. Sampling was stratified by sex, reflecting the primary comparison of interest in this manuscript. No formal 1:1 matching procedure was applied. Instead, participants were drawn using stratified random sampling, with age in months used as the stratification variable to ensure that the male and female subsamples had comparable age distributions across the full 107-132 month range. Because the ABCD baseline sample is restricted to children aged 9-11 years, the age range is inherently narrow, making strict individual-level matching unnecessary to achieve comparability. The female subsample consisted of 245 participants aged 107-132 months (mean = 119 months; SD = 7.74 months). The male subsample consisted of 255 participants aged 107-132 months (mean = 119.18 months; SD = 7.36 months). Demographic and scanner-related characteristics, including site distribution, are summarized in Table 1.

**Table 1.** Overview of demographic and scanner-related characteristics by sex. Sex ratio and age p-values confirm a balanced group composition. IQ scores reflect the uncorrected NIH Toolbox total composite scores. Cognitive measures include NIH Toolbox Cognition Battery tasks and the Rey Auditory Verbal Learning Test (RAVLT); uncorrected scores reflect raw performance, age-corrected scores are normed relative to age-matched peers, and T-scores are standardized to a normative sample mean of 50 (SD = 10). MRI scanner site reflects the scanner manufacturer used for data acquisition (Siemens, Philips, or GE). ABCD site reflects the data collection site within the Adolescent Brain Cognitive Development (ABCD) study. Parental education represents the highest level of education reported by either parent, with “college degree or higher” defined as completion of a bachelor’s degree, master’s degree, professional degree, or doctoral degree. Parental employment indicates whether at least one parent was employed full-time. Household income was grouped into three categories, low (<$35,000), middle ($35,000-$99,999), and high (≥$100,000). Race/ethnicity was reported by parents and categorized as White, Black or African American or Other (including Hispanic, Asian, and multiracial individuals). P-values indicate the statistical significance of differences between males and females for each category. P-values are uncorrected for multiple comparisons and are reported descriptively to characterize sample balance. Only three comparisons (all RAVLT scores) had uncorrected p < 0.05, and none remained significant after false discovery rate (FDR) correction across all comparisons in the table (minimum q = 0.12).

| **Variables** | **Female** | **Male** | **p value** |
|---|---|---|---|
| **N** | 245 | 255 | - |
| **Sex Ratio** | 49% | 51% | 0.65 |
| **Age (mean ± SD)** | 119 ± 7.74 | 119.18 ± 7.36 | 0.34 |
| **IQ (mean ± SD)** | 85.86 ± 13.07 | 85.50 ± 10.35 | 0.15 |
| **Flanker Inhibitory Control and Attention:** |  |  |  |
| Uncorrected (mean ± SD) | 93.30 ± 9.15 | 94.50 ± 8.83 | 0.15 |
| Age-corrected scores (mean ± SD) | 94.41 ± 13.92 | 96.10 ± 13.72 | 0.19 |
| **List Sorting Working Memory:** |  |  |  |
| Uncorrected (mean ± SD) | 96.86 ± 12.46 | 97.27 ± 13.48 | 0.73 |
| Age-corrected scores (mean ± SD) | 101.36 ± 16.65 | 101.35 ± 16.88 | 0.99 |
| **Dimensional Change Card Sort:** |  |  |  |
| Uncorrected (mean ± SD) | 93.06 ± 9.60 | 92.77 ± 8.97 | 0.77 |
| Age-corrected scores (mean ± SD) | 98.30 ± 15.85 | 96.75 ± 15.91 | 0.29 |
| **Pattern Comparison Processing Speed:** |  |  |  |
| Uncorrected (mean ± SD) | 89.72 ± 14.68 | 88.33 ± 14.68 | 0.31 |
| Age-corrected scores (mean ± SD) | 96.52 ± 22.88 | 93.86 ± 22.41 | 0.21 |
| **Picture Vocabulary:** |  |  |  |
| Uncorrected (mean ± SD) | 84.43 ± 8.21 | 84.97 ± 7.75 | 0.46 |
| Age-corrected scores (mean ± SD) | 107.11 ± 17.58 | 107.80 ± 16.60 | 0.66 |
| T-score (mean ± SD) | 0.47 ± 1.72 | 0.58 ± 1.62 | 0.45 |
| **Picture Sequence Memory:** |  |  |  |
| Uncorrected (mean ± SD) | 103.39 ± 11.61 | 101.74 ± 11.60 | 0.12 |
| Age-corrected scores (mean ± SD) | 101.78 ± 15.47 | 99.56 ± 15.35 | 0.12 |
| T-score (mean ± SD) | -0.47 ± 0.73 | -0.58 ± 0.73 | 0.12 |
| **Oral Reading Recognition:** |  |  |  |
| Uncorrected (mean ± SD) | 90.80 ± 6.95 | 91.39 ± 6.55 | 0.35 |
| Age-corrected scores (mean ± SD) | 102.48 ± 19.52 | 103.83 ± 18.10 | 0.44 |
| T-score (mean ± SD) | -0.39 ± 2.40 | -0.17 ± 2.27 | 0.32 |
| **Fluid Cognition Composite:** |  |  |  |
| Uncorrected (mean ± SD) | 92.18 ± 10.83 | 91.40 ± 11.89 | 0.46 |
| Age-corrected scores (mean ± SD) | 95.84 ± 21.76 | 94.09 ± 21.40 | 0.38 |
| **Crystallized Cognition Composite:** |  |  |  |
| Uncorrected (mean ± SD) | 85.92 ± 9.06 | 86.96 ± 6.71 | 0.16 |
| Age-corrected scores (mean ± SD) | 103.92 ± 23.23 | 105.73 ± 21.10 | 0.38 |
| **Total Cognition Composite:** |  |  |  |
| Uncorrected (mean ± SD) | 86.17 ± 10.82 | 86.33 ± 10.41 | 0.86 |
| Age-corrected scores (mean ± SD) | 99.68 ± 23.33 | 99.66 ± 21.80 | 0.99 |
| **RAVLT:** |  |  |  |
| Immediate Recall T-score (mean ± SD) | 4.87 ± 1.67 | 4.88 ± 1.85 | 0.92 |
| Delayed Recall T-score (mean ± SD) | 9.99 ± 2.84 | 9.22 ± 3.31 | 0.008 |
| Recognition Hits T-score (mean ± SD) | 9.59 ± 3.00 | 8.85 ± 3.30 | 0.01 |
| Semantic/Category Recall T-score (mean ± SD) | 45.88 ± 9.67 | 43.64 ± 9.95 | 0.01 |
| **MRI Scanner Site** | 60.3% Siemens<br>14.8% Philips<br>24.8% GE | 62.0% Siemens<br>12.4% Philips<br>25.6% GE | 0.52 |
| **ABCD Site** | 5.5% UCLA<br>6.2% UC San Diego<br>2.9% SRI International<br>5.8% FIU<br>4.8% MUSC<br>4.5% OHSU<br>5.4% University of Utah<br>2.8% CU Boulder<br>3.3% LIBR<br>2.9% Washington University<br>4.1% University of Minnesota<br>3.3% UWM<br>7.0% U-M<br>5.4% UVM<br>4.1% UR<br>9.5% Yale University<br>6.9% University of Pittsburgh<br>5.3% University of Maryland<br>3.7% VCU<br>4.1% UF<br>2.5% CHLA | 1.9% UCLA<br>4.3% UC San Diego<br>5.8% SRI International<br>5.8% FIU<br>2.3% MUSC<br>6.2% OHSU<br>3.5% University of Utah<br>4.3% CU Boulder<br>3.1% LIBR<br>7.0% Washington University<br>6.0% University of Minnesota<br>4.7% UWM<br>5.9% U-M<br>9.3% UVM<br>4.3% UR<br>3.1% Yale University<br>6.2% University of Pittsburgh<br>5.0% University of Maryland<br>4.7% VCU<br>4.1% UF<br>2.5% CHLA | 0.71 |
| **Parental Education** | 18.6% high school degree or less<br>50.0% college degree or higher | 18.5% high school degree or less<br>59.3% college degree or higher | 0.14 |
| **Parental Employment** | 73.5% full-time employed | 74.8% full-time employed | 0.37 |
| **Household Income** | 19.4% low<br>36.4% middle<br>38.8% high | 22.5% low<br>26.7% middle<br>45.3% high | 0.58 |
| **Race/Ethnicity** | 57.0% White<br>9.1% Black or African American<br>33.9% other | 48.4% White<br>12.4% Black or African American<br>39.2% other | 0.06 |

### B. MRI Parameters and Preprocessing

Data were collected on three types of 3T scanners (Siemens Prisma, General Electric 750 and Philips) all with a standard adult-size 32-channel head coil. The MRI sequences acquired and used for this work were a T1 structural scan (TR = 2500/2500/6.31 ms respectively; TE = 2.88/2/2.9 ms respectively). The resting-state EPI sequence used the following parameters: TR = 800 ms, TE = 30 ms, 60 slices, flip angle = 52^°^, matrix size = 90 × 90, FOV = 216 × 216 mm, resolution = 2.4 × 2.4 × 2.4 mm. The full details of the imaging acquisition protocol are described in (Casey et al., 2018).

Structural MRI data were preprocessed using the Statistical Parametric Mapping (SPM) software. Spatial normalization was performed using the DARTEL (Diffeomorphic Anatomical Registration Through Exponentiated Lie Algebra) algorithm to create a study-specific template and register images to MNI space with high precision. The resulting gray matter maps were modulated to preserve total tissue volume and resampled to a voxel size of 1.5 × 1.5 × 1.5 mm. Finally, the maps were smoothed with a 4 mm full width at half-maximum (FWHM) Gaussian kernel to reduce noise while preserving local anatomical detail.

The data preprocessing of the resting-state sequence involved a combination of the FMRIB Software Library v6.0 (FSL) toolbox and the Statistical Parametric Mapping 12 (SPM) toolbox in MATLAB 2020b. To correct for subject head motion, the FSL MCFLIRT tool was employed to perform rigid body motion correction. Following the motion correction, the distortion in the fMRI images was addressed using field map files. These field maps were obtained by capturing volumes with phase encoding in the anterior-posterior (AP) and posterior-anterior (PA) directions using the FSL tool topup. By utilizing the output field map coefficients obtained from the FSL tool applytopup, the distortion in the fMRI volume was corrected. Subsequently, the fMRI data were smoothed using a Gaussian kernel with a full width at half maximum (FWHM) of 6 mm and warped to the standard Montreal Neurological Institute (MNI) space with a spatial resolution of 3 × 3 × 3 mm. Following spatial preprocessing, 53 ICN timecourses were extracted using the NeuroMark_fMRI_1.0 constrained ICA pipeline implemented in the GIFT toolbox. Timecourses were subsequently denoised by removing linear trends and suppressing outlier spikes, bandpass filtered using a 7th-order Butterworth filter (0.01–0.15 Hz), and z-score normalized prior to analysis.

### C. Dynamic Inter-Modality Source Coupling

We used a standardized functional template consisting of multiple replicable resting fMRI brain networks, estimated via ICA (Lee, 1998), as spatial priors for both structural and functional modalities. This approach ensured consistency and comparability across subjects while also adapting to each subject individually. Specifically, we used fully automated spatially constrained ICA (scICA), applied separately to rs-fMRI and sMRI data, to extract subject-specific components while preserving spatial independence (Kotoski et al., 2024). This enabled us to evaluate cross- modal coupling within a unified framework, where rs-fMRI captures individual dynamic patterns and structural brain morphometry reflects inter-subject covariation. We used the 53-component NeuroMark_fMRI_1.0 template (Du et al., 2020) as the spatial prior, ensuring that the resulting components were both data-adaptive and comparable across modalities. The template organizes the 53 components into 7 functional domains: subcortical, auditory, sensorimotor, visual, cognitive-control, default-mode, and cerebellar. A full description of the brain regions comprising each domain is provided in Figure 1. Although the NeuroMark template was originally developed on functional data, it has been successfully applied to structural data in prior work (Kotoski et al., 2024), validating its use as a spatial prior for SBM component extraction. All analyses were conducted using the GIFT toolbox (Calhoun et al., 2001), an open-source MATLAB-based software package for performing group-level and subject-specific independent component analysis on neuroimaging data. GIFT implements spatially constrained ICA, which allows subject-specific component estimation by using a group-level spatial template as a prior, ensuring that the resulting components are both data-adaptive and comparable across subjects. It was selected for the present study because it natively supports constrained ICA for both rs-fMRI and sMRI data within a unified framework, enabling consistent component extraction across modalities and facilitating the cross-modal coupling analysis central to dIMSC.

**Figure 1.**
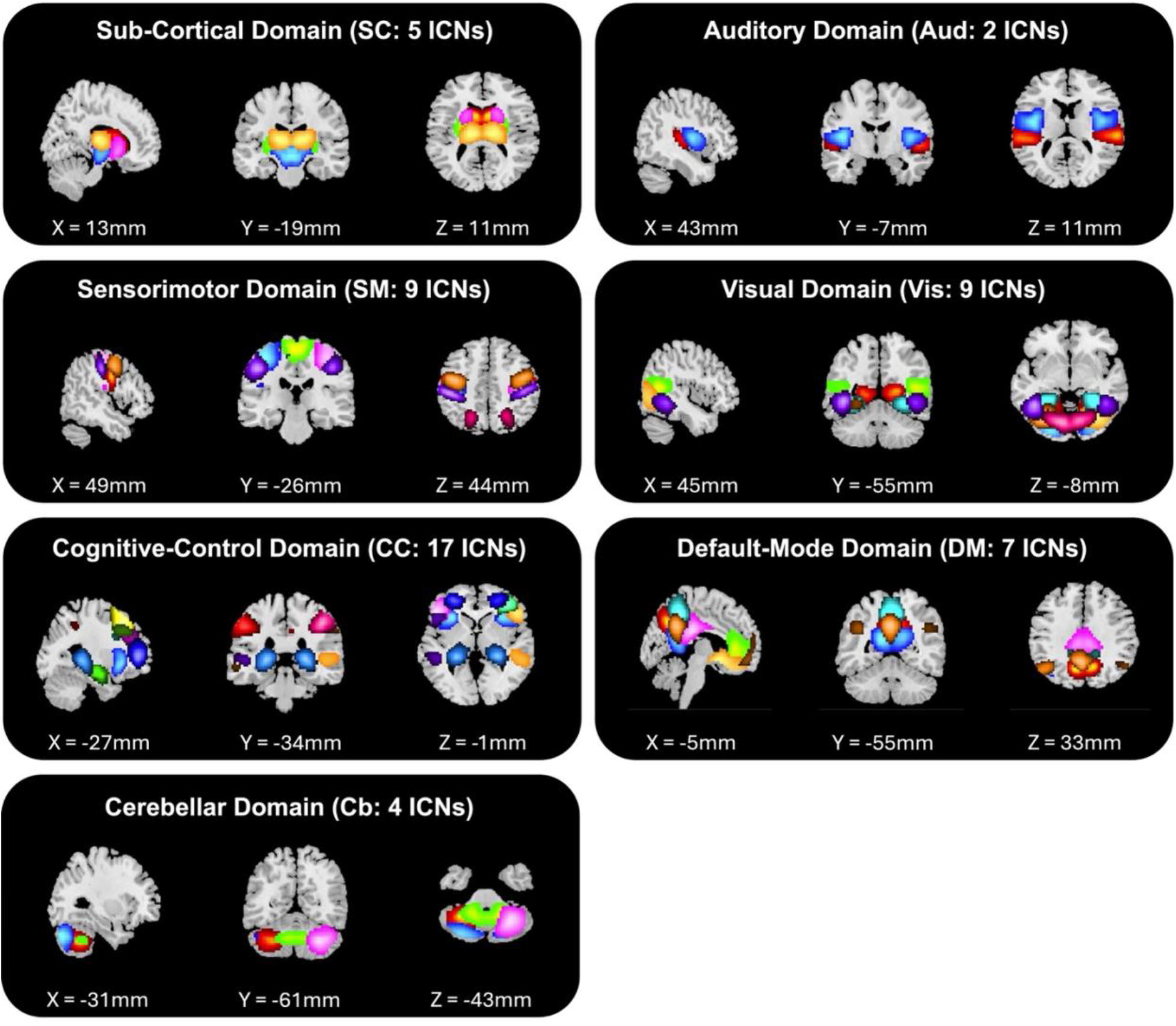
The NeuroMark_fMRI_1.0 functional template organized into 7 functional domains. The template comprises 53 intrinsic connectivity networks (ICNs) grouped into 7 functional domains: subcortical (SC: 5 ICNs), auditory (Aud: 2 ICNs), sensorimotor (SM: 9 ICNs), visual (Vis: 9 ICNs), cognitive-control (CC: 17 ICNs), default-mode (DM: 7 ICNs), and cerebellar (Cb: 4 ICNs). Brain maps are displayed in MNI space with representative axial, coronal, and sagittal slices for each domain. The template was applied as a spatial prior for constrained independent component analysis of both resting-state fMRI and structural MRI data, ensuring consistent and comparable component extraction across subjects and modalities. Adapted from Du et al. (2020).

For rs-fMRI data, we computed a dynamic functional network connectivity matrix using sliding window Pearson correlation (SWPC) (Hindriks et al., 2016), which estimates time-varying functional interactions. The analysis employed a rectangular window of 44s with a step size of 1 TR (Allen et al., 2014; Wiafe et al., 2024), yielding 304 windows, allowing us to capture fluctuations in connectivity over time. This approach provides a dynamic view of functional organization beyond static connectivity measures by characterizing how interactions between brain networks evolve temporally. Note that throughout this manuscript, “timepoints” refers to these 304 sliding windows rather than individual TRs, each window represents a snapshot of functional connectivity computed over a 44s interval, stepped by one TR across the full scan duration.

For sMRI analysis, we applied constrained SBM to estimate subject-specific loading parameters for 53 brain regions (Kotoski et al., 2024). SBM is a data-driven multivariate approach that identifies spatially independent structural patterns and quantifies their expression in individual subjects (Gupta et al., 2019). Using gray matter volume as input, we extracted independent components that capture covarying patterns across subjects. Each subject’s structural profile was represented by a 53-component loading vector, where each component reflects the degree to which the corresponding gray matter pattern is expressed in that individual. SBM loading parameters were estimated from all 500 participants simultaneously, regardless of sex. This approach was deliberately chosen to ensure that the structural components are anchored to the same spatial reference for both males and females, preserving cross-group comparability of the resulting loading vectors. Estimating components separately by sex would risk introducing group-specific spatial solutions that are not directly comparable, undermining the validity of subsequent sex difference analyses.

To link the two modalities within our proposed dIMSC approach, we estimate the network expression coupling between the dFNC pattern and the SBM pattens via cross-correlation, computed at each time step *w* across 304 sliding windows between the dFNC matrix and the corresponding SBM vector for each subject:

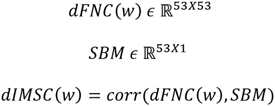

Where:

- *dIMSC*(*w*) *ε* ℝ^53*X*1^is the dynamic inter-modality source coupling at window w for each subject.
- *dFNC*(*w*) *ε* ℝ^53*X*53^ is the dynamic functional connectivity matrix at window w for each subject.
- *SBM ε* ℝ^53*X*1^ is the structural component vector for each subject.

This correlation results in a timecourse that indicates how the functional characteristics of the network (represented as a column in the dFNC matrix) align with the expression of covarying gray matter components (represented as a row in the SBM mixing matrix) in each subject. This analysis resulted in a matrix that showed the degree of dynamic structure-function coupling. Figure 2 shows the pipeline for the dIMSC method. Subsequently, we categorized each timepoint based on a correlation threshold for each of the 53 brain networks based on the distribution percentiles of the dIMSC values. We selected a threshold of *ρ* = ±0.1, such that approximately 50% of the timepoints fall into a neutral state, while the positive and negative states each contain approximately 25%. This distribution aligns with the expected behavior of coupling, where fluctuations around a stable mean are more common, and larger deviations occur less frequently (Hutchison et al., 2013). This ensures that the positive and negative states reflect meaningful changes in coupling while maintaining balanced subgroup sizes. On average, across all participants, 27.12% of timepoints exhibited positive coupling, 46.62% were classified as neutral, and 26.26% displayed negative coupling. For each category of the degree of time-resolved (or dynamic) structure-function coupling values, we computed the occupancy rate, representing the proportion of time a network spends in a particular coupling state.

**Figure 2.**
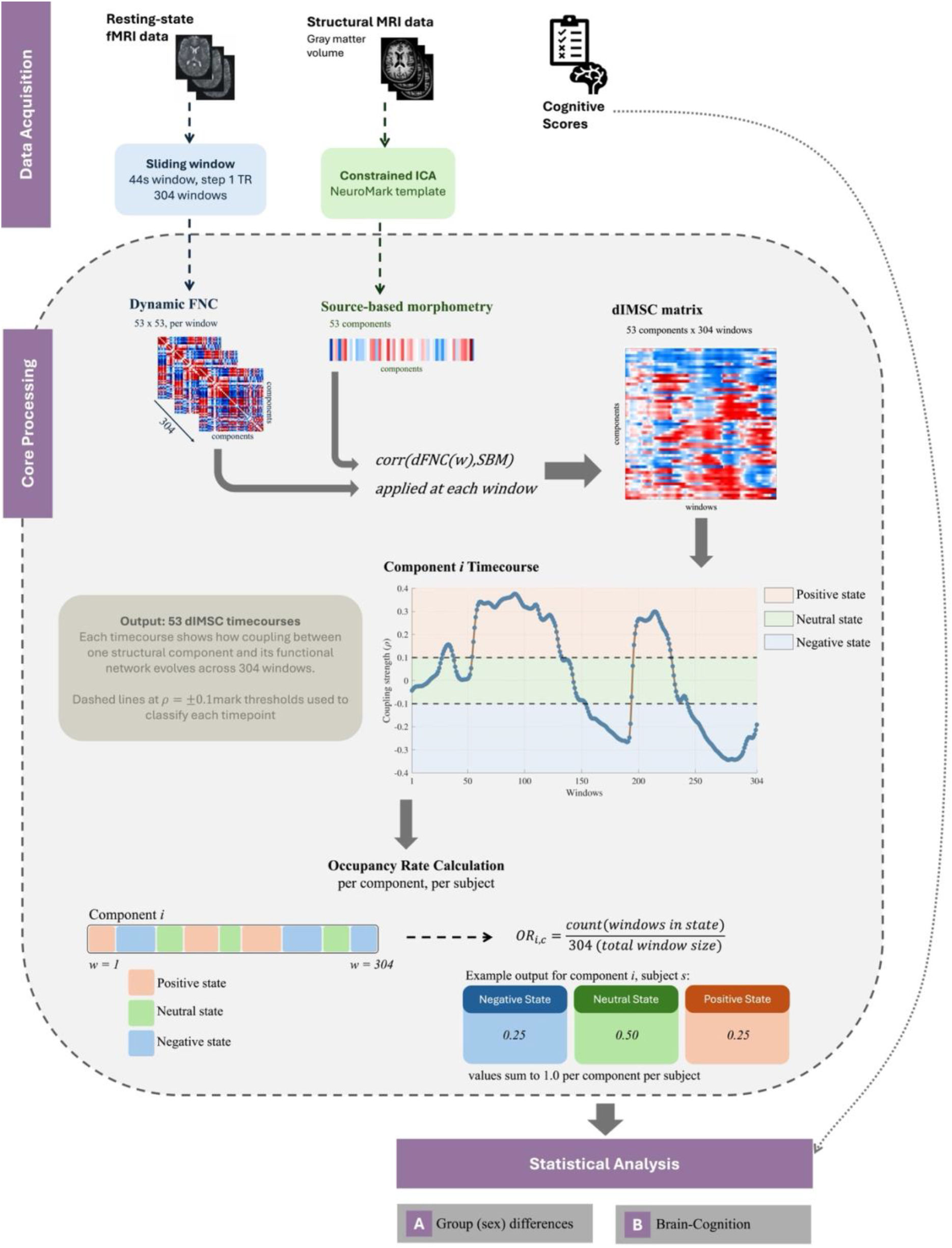
Overview of the dIMSC pipeline. Top panel (Data Acquisition): resting-state fMRI data are processed using a sliding window approach (44s window, step 1 TR, yielding 304 windows) to compute dynamic functional network connectivity (dFNC) matrices (53 × 53, per window). Structural MRI gray matter volume is processed using constrained ICA with the NeuroMark_fMRI_1.0 template to generate source-based morphometry (SBM) loading vectors (53 components). Cognitive scores are included for the moderation analysis. Core Processing panel: at each window w, dIMSC is computed as the cross-correlation between the dFNC matrix and the SBM vector, yielding a dIMSC matrix (53 components × 304 windows). Each row of this matrix represents a component timecourse reflecting the time-resolved coupling strength between structural and functional patterns. Timepoints are classified into three coupling states based on the empirical distribution of dIMSC values: positive state (ρ > 0.1, ∼25% of timepoints), neutral state (−0.1 ≤ ρ ≤ 0.1, ∼50% of timepoints), and negative state (ρ < −0.1, ∼25% of timepoints). The occupancy rate (OR) is then computed as the proportion of windows spent in each state per component per subject, yielding three values per component that sum to 1.0. Bottom panel (Statistical Analysis): occupancy rates serve as input to two analyses: (A) sex group differences across 53 components and (B) brain-cognition moderation analysis.

The neutral state correlation values near zero represents timepoints where the dynamic inter-modality structure-function coupling is weak. Importantly, this state should not be interpreted as a distinct or meaningful coupling state, but rather as reflecting a stable period without significant alignment between structural and functional patterns. It serves as a reference against which deviations towards stronger positive or negative coupling can be identified or characterized. By distinguishing these neutral timepoints, we can better capture meaningful fluctuations in coupling dynamics while accounting for period of relative stability.

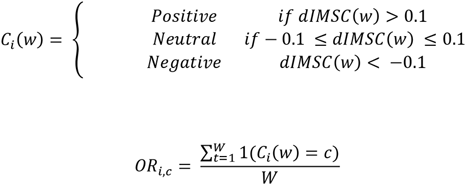

where:

- *OR_i_*_,*c*_ is the occupancy rate for state *c* in brain network *i*.
- *W* is the total number of windows.
- 1(*C_i_*(*w*) = *c*) is an indicator function that equals 1 if *C_i_*(*w*) is in state *c* and 0 otherwise.

Key terminology used throughout this manuscript is defined in the Technical Terms document and, with additional methodological detail, in Supplementary Table 1.

To examine sex differences in dIMSC measures, we conducted a multiple linear regression including a regressor for age (in months), site, and motion (mean framewise displacement), intracranial volume computed using FreeSurfer v6.0 (Fischl, 2012), as a nuisance regressor. A significance threshold of *p* < .05 was used, with corrections for multiple comparisons applied using the false discovery rate (FDR).

### D. Sex Moderation of dIMSC-Behavior Relationships

To validate the functional sensitivity of the dIMSC framework, we examined whether the coupling states identified by dIMSC carry neurobiological meaning by testing whether they differentially predict cognitive performance as a function of biological sex. This moderation analysis serves as a proof-of-concept for the method: if dIMSC coupling states capture meaningful variation in brain organization, they should be systematically related to cognition in a sex-dependent manner consistent with known neurodevelopmental patterns. The objective was to determine if biological sex significantly modulates the relationship between the dIMSC occupancy rate and a set of cognitive and behavioral measures. The analysis involved testing the relationship between the state-specific occupancy rate of the 53 brain networks and a total of 27 cognitive/behavioral measures across all three coupling states (positive, neutral, negative):

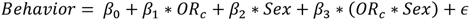

Where *OR_c_* is the occupancy rate in state *c* (positive, neutral, negative). Sex is a binary value (male or female), and *β*_3_ is the coefficient for the interaction term (*OR_c_* ∗ *Sex*), indicating that biological sex moderated the relationship. Occupancy rates and cognitive scores were z-scored before model fitting, so that β₃ is a standardized coefficient comparable across measures with different scales; standardization does not change the associated p-values. The model also included the following covariates: age (in months), site, intracranial volume, and motion (mean framewise displacement), and all remaining neurocognitive scores as mutual confounds. A significance threshold of p < 0.05 was used, with corrections for multiple comparisons applied across all 53 networks, 3 coupling states, and 27 behavioral measures using false discovery rate.

We utilized a total of 27 measures spanning executive function, language, reading, fluid/crystallized cognition, and memory from the NIH Toolbox and the Rey Auditory Verbal Learning Test (RAVLT), which are administered as part of the standard ABCD study protocol and have been widely used to characterize cognitive sex differences in pediatric samples. The specific measures included in the moderation analysis were:

1. NIH Toolbox Executive Function:

- Flanker Inhibitory Control and Attention: Uncorrected and age-corrected scores
- List Sorting Working Memory: Uncorrected and age-corrected scores
- Dimensional Change Card Sort: Uncorrected and age-corrected scores
- Pattern Comparison Processing Speed: Uncorrected and age-corrected scores

2. NIH Toolbox Language & Reading:

- Picture Vocabulary: Uncorrected score, age-corrected score, and T-score
- Picture Sequence Memory: Uncorrected score, age-corrected score, and T-score
- Oral Reading Recognition: Uncorrected score, age-corrected score, and T-score

3. NIH Toolbox Composite Scores:

- Fluid Cognition Composite: Uncorrected and age-corrected scores
- Crystallized Cognition Composite: Uncorrected and age-corrected scores
- Total Cognition Composite: Uncorrected and age-corrected scores
- RAVLT Memory Measures:
- RAVLT Immediate Recall (Trial I-V Sum): T-score
- RAVLT Delayed Recall (Trial VII): T-score
- RAVLT Recognition Hits (Trial VI): T-score
- RAVLT Semantic/Category Recall (List B Interference): T-score

For measures where multiple score types were available, we included uncorrected scores, age-corrected scores, and T-scores separately. Uncorrected scores reflect raw task performance; age-corrected scores remove the influence of age-related variance, isolating individual differences in performance relative to age-matched peers; and T-scores provide standardized values relative to a normative sample. Including all three score types allowed us to examine whether the structure-function coupling relationships with cognition were robust across different ways of representing performance, and in particular whether they persisted after accounting for age effects.

## III. RESULTS

The application of dIMSC revealed distinct temporal patterns of structure-function coupling across the 53 brain networks. By estimating the coupling between the network expression of the dFNC matrix and the SBM vector at each timepoint, we obtained a time-resolved measure of coupling strength. These values were grouped into three states: positive, neutral, and negative coupling. Specifically, positive coupling indicates that increases in structural properties are associated with increases in functional properties, suggesting a coordinated relationship; neutral coupling indicates that structural and functional properties are relatively independent of each other; and negative coupling implies an inverse relationship, potentially indicating compensatory or opposing mechanisms between the areas.

The mean coupling strength was 0.18 ± 0.04 for positive states, 0.0001 ± 0.02 for neutral states, and -0.17 ± 0.04 for negative states. This distribution indicates a predominant trend of balanced structure-function alignment across the sample, with stronger positive and negative coupling suggesting a bidirectional relationship between structural and functional features in specific networks. When analyzed separately by sex, males exhibited positive coupling in 26.96% of timepoints, neutral coupling in 47.16%, and negative coupling in 25.89%. The corresponding mean coupling strengths were 0.18 ± 0.04 for positive states, 0.0002 ± 0.02 for neutral states, and -0.17 ± 0.04 for negative states. In females, positive coupling was observed in 27.30% of timepoints, neutral coupling in 46.08%, and negative coupling in 26.63%, with mean coupling strengths of 0.17 ± 0.04, 5.43 × 10⁻⁶ ± 0.02, and -0.17 ± 0.04, respectively. Descriptively, females showed a slightly greater proportion of timepoints classified into the positive and negative coupling states relative to males, though this difference was not formally tested. On average, males had a total intracranial volume of 1583.55 ± 119.96 cm³, while females had an average intracranial volume of 1438.48 ± 191.21 cm³. To account for potential head size effects, all statistical comparisons were adjusted for intracranial volume.

Sex differences in dIMSC were observed across several brain networks. Figure 3 provides spatial brain maps highlighting the identified regions, while Figure 4 presents the occupancy rate values for significant brain regions in both males and females. To provide a transparent overview of sex differences in dIMSC occupancy rate across all 53 components and three coupling states prior to the brain map visualization, FDR-corrected p-values for group comparisons are reported in Supplementary Table 2. Within the positive coupling state, females exhibited stronger coupling in the inferior parietal lobule and precuneus compared to males, suggesting enhanced structural support for functional activity in these regions. In the neutral coupling state, males demonstrated stronger coupling in the cerebellum and superior temporal gyrus, while females exhibited stronger coupling in the caudate, precentral, and middle frontal gyrus, suggesting that different brain regions may underlie sex-specific structure-function coupling during this state. For the negative coupling state, males showed stronger coupling in the paracentral lobule, superior parietal lobule, and middle temporal gyrus, while females exhibited stronger coupling in the superior temporal gyrus and inferior parietal lobule.

**Figure 3.**
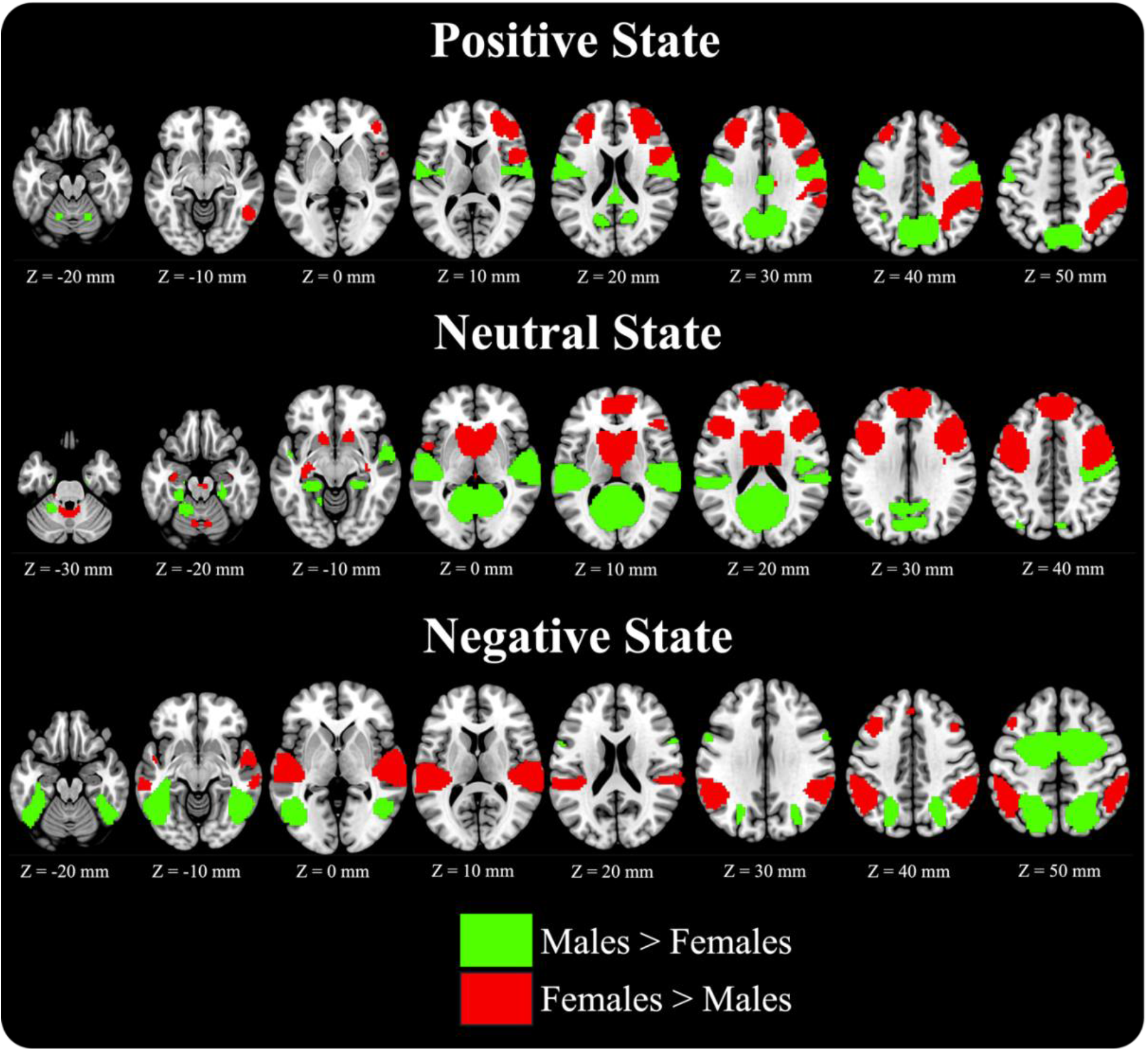
Sex Differences in Dynamic Structure-Function Coupling Occupancy Rate Across 53 Brain Components. Brain maps display components with statistically significant sex differences in dIMSC occupancy rate, shown separately for the positive state (top, ρ > 0.1), neutral state (middle, −0.1 ≤ ρ ≤ 0.1), and negative state (bottom, ρ < −0.1). Components are color-coded categorically by direction of effect: red indicates components where females show significantly greater occupancy (F > M) and green indicates components where males show significantly greater occupancy (M > F). Non-significant components are not shown. Statistical significance was determined using two-sample t-tests with FDR correction across all 53 components and 3 coupling states (p < 0.05). Anatomical localization is shown in MNI space.

**Figure 4.**
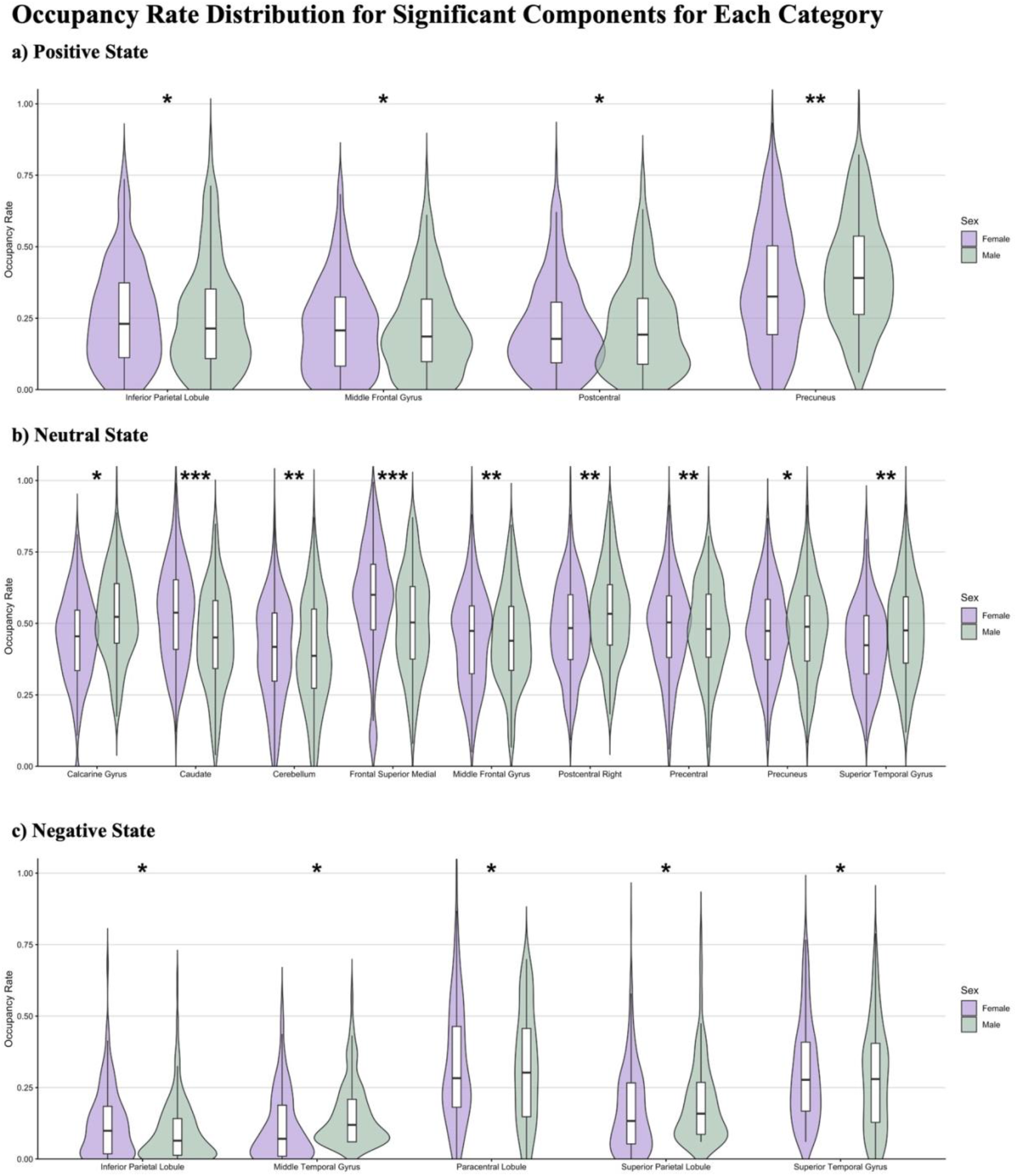
State-level violin plots displaying state-specific occupancy rates for brain regions that exhibited significant sex differences, with values shown separately for males (green) and females (purple). The y-axis represents the occupancy rate, defined as the proportion of timepoints during which each brain region was classified within a specific coupling state, ranging from 0 to 1. Note that the y-axis reflects occupancy rate values (0-1), not dIMSC correlation values; the ±0.1 thresholds are used upstream to classify each timepoint into a coupling state, and the occupancy rate is computed subsequently as a summary proportion. The x-axis lists the individual brain regions for each group. Asterisks indicate statistically significant sex differences (* for 0.05 > p > 0.01, ** for 0.01 > p > 0.001, and *** for p < 0 .001), corrected for multiple comparisons using FDR. Notably, sex differences appear relatively evenly distributed across all three coupling states. This suggests that sex differences in brain activity patterns occur throughout the full range of brain network dynamics. This pattern might reflect that biological sex influences the coordinated brain activity in general, regardless of the specific coupling strength.

To validate the functional sensitivity of dIMSC, we examined whether the coupling states identified by the method differentially predict cognitive performance as a function of biological sex. The moderation analysis revealed significant sex-dependent associations between dIMSC coupling states and 27 cognitive and behavioral measures across all three coupling states (positive, neutral, and negative). These results confirm that the time a brain network spends in a specific dIMSC state carries neurobiological meaning, as its relationship to cognitive performance is systematically modulated by sex. The beta coefficients for the significant interaction terms are summarized in the heatmaps (Figure 5).

**Figure 5.**
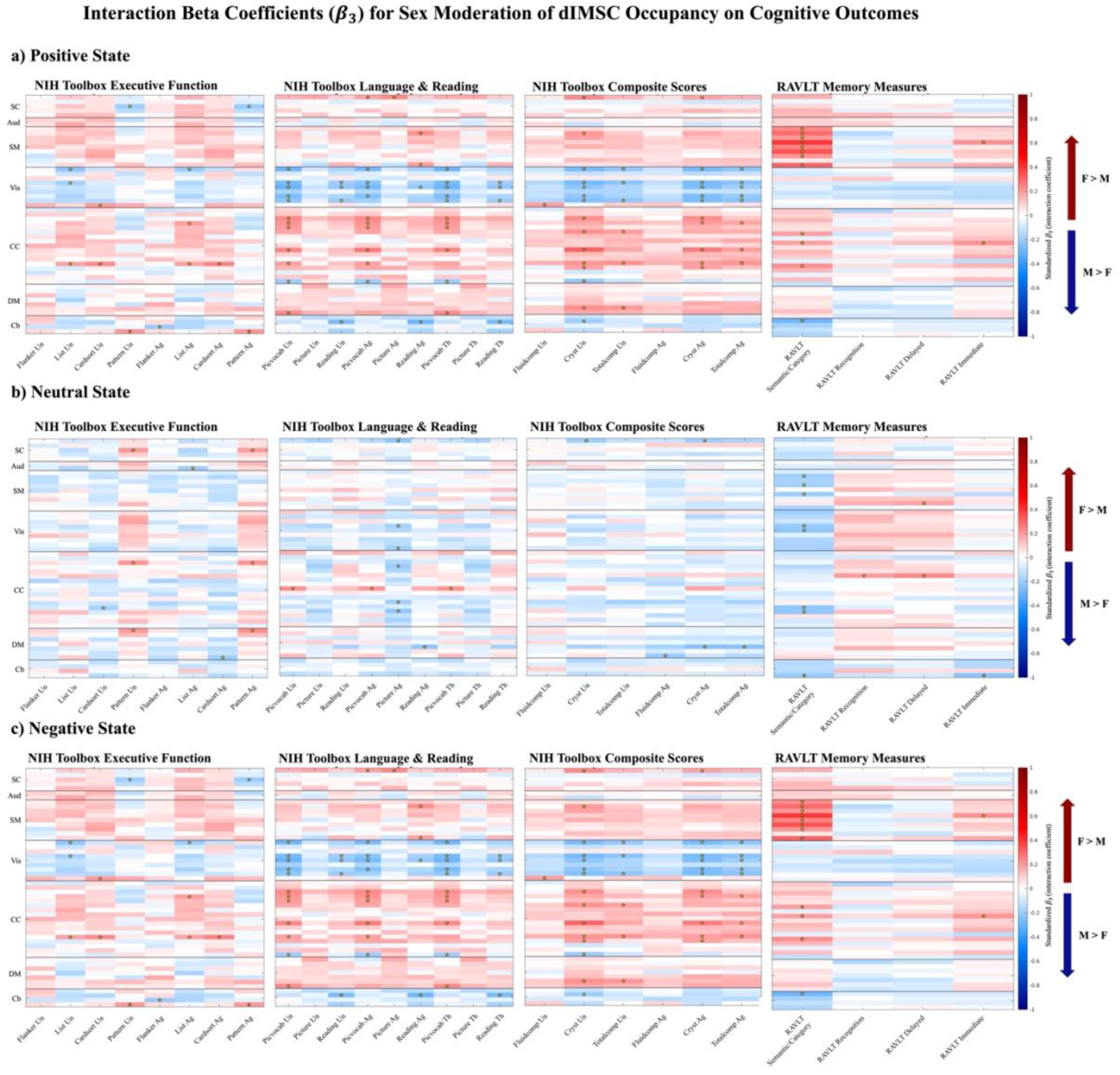
This figure displays the standardized *β*_3_ coefficients from the moderation analysis. Panels correspond to the three coupling states: (a) positive state (ρ > 0.1), (b) neutral state (−0.1 ≤ ρ ≤ 0.1), and (c) negative state (ρ < −0.1). The y-axis represents the 53 brain networks components, grouped by domains (SC: subcortical, Aud: auditory, SM: sensorimotor, Vis: Visual, CC: cognitive-control, DM: default-mode, Cb: cerebellar). The x-axis lists the 27 cognitive and behavioral measures grouped by domain: NIH Toolbox Executive Function, NIH Toolbox Language & Reading, NIH Toolbox Composite Scores, and RAVLT Memory Measures. Color reflects the direction and magnitude of the interaction effect: red (positive β₃) indicates that the association between dIMSC occupancy and cognitive performance is stronger in females than males (F > M); blue (negative β₃) indicates that the association is stronger in males than females (M > F). Stars denote statistically significant interactions after FDR correction across all 53 components, 3 coupling states, and 27 behavioral measures (p < 0.05). This figure serves as proof-of-concept validation for dIMSC, demonstrating that the coupling states it identifies are neurobiologically meaningful and differentially predictive of cognitive performance as a function of biological sex. Coefficients are standardized (occupancy rate and cognitive score were z-scored) and the color scale is fixed to [−1, 1]. Abbreviations: Un, uncorrected score; Ag, age-corrected score; Th, T-score; Flanker, Flanker Inhibitory Control and Attention; List, List Sorting Working Memory; Cardsort, Dimensional Change Card Sort; Pattern, Pattern Comparison Processing Speed; Picvocab, Picture Vocabulary; Picture, Picture Sequence Memory; Reading, Oral Reading Recognition; Fluidcomp, Fluid Cognition Composite; Cryst, Crystallized Cognition Composite; Totalcomp, Total Cognition Composite; RAVLT Semantic/Category, Semantic/Category Recall (List B interference); RAVLT Recognition, Recognition Hits (Trial VI); RAVLT Delayed, Delayed Recall (Trial VII); RAVLT Immediate, Immediate Recall (Trial I–V sum); all RAVLT scores are T-scores.

The positive coupling state, which reflects strong structural-functional alignment, exhibited significant and distinct sex-specific associations with cognition. We observed female-dominant interactions where higher occupancy in the positive state was more strongly associated with better performance in females, particularly for crystallized cognition measures. These interactions were centered in regions including the thalamus, calcarine sulcus, cerebellum, and other areas of the visual cortex. Conversely, males exhibited stronger reliance on the positive coupling state for better performance across measures of executive function and working memory. These male-dominant interactions involved the thalamus, cerebellum, calcarine sulcus, hippocampus, postcentral, and paracentral gyri. The thalamus emerged as a central hub for interactions across multiple measures.

The neutral coupling state, characterized by weak structure-function alignment, also demonstrated significant moderation effects, with almost all interactions for working memory and fluid reasoning being associated with males. For executive function, interactions were mixed; while some were evident in males, such as the card sort test, interactions related to pattern comparison processing speed were generally more pronounced in females. The prevalent M>F pattern for working memory and fluid reasoning suggests that males exhibit a more widespread dependence on certain areas to predict their cognitive performance, even when the dIMSC coupling is weak. This may reflect a reliance on more robust neural networks in males when structural-functional alignment is relatively low.

The negative coupling state, representing an inverse relationship between structure and function, showed strong, widespread interactions, often stronger in females. Significant interactions in females were observed across multiple domains, including working memory and fluid reasoning, involving sensorimotor and attentional networks such as the postcentral and paracentral gyri, inferior parietal lobule, cerebellum, and superior parietal lobule. Strong female interactions were also found in language-related areas and memory/visual areas for verbal and crystallized cognition, while stronger male interactions were present in frontal superior areas.

## IV. DISCUSSION

In this study, we introduced dynamic inter-modality source coupling, a novel method to quantify the time-varying constraints that brain structure imposes on functional dynamics.

Extending our previous IMSC method (Kotoski et al., 2024), dIMSC moves beyond the assumption that structure-function relationships are constant, instead modeling them as fluctuating states of alignment (positive), independence (neutral), or divergence (negative). By applying this framework to a large sample of children from the ABCD study, we demonstrate that the coupling between gray matter architecture and functional connectivity is not uniform but state-dependent, and the organization of these states exhibits significant sex differences that are relevant to cognitive performance.

The classification of coupling into positive, neutral, and negative states provides a theoretical framework for interpreting neurodevelopmental dynamics. Before interpreting directionality, we clarify what “expression” means in this context: SBM expression is a subject-specific loading coefficient obtained from spatially-constrained ICA of gray matter volume maps, reflecting how strongly an individual’s gray matter pattern matches a given structural covariance component; it is a continuous multivariate score of structural covariance, not a direct measure of tissue volume or thickness at a single location. Throughout this section, references to a region’s “structural substrate” or “structural scaffold” refer to this SBM loading. Positive coupling reflects periods where functional connectivity patterns tightly align with the underlying structural scaffold, networks with greater gray matter expression show proportionally stronger functional connectivity at that timepoint. This is consistent with the “structural constraint” hypothesis where the anatomical architecture of a region directly limits or shapes the range of functional dynamics it can express (Baum et al., 2020; Honey et al., 2009). In other words, when a region has a strong structural substrate, its functional activity during those timepoints is being actively shaped or ’pulled’ toward that substrate. Negative coupling, by contrast, represents an inverse relationship, where regions with greater gray matter expression show relatively weaker functional connectivity at that timepoint. Rather than reflecting structural constraints, this state is more consistent with active functional divergence or segregation, periods where functional networks temporarily decouple from their structural basis, potentially to support flexible, transient cognitive operations that require dynamics beyond what the structural scaffold would ordinarily permit (Wan et al., 2022). The key distinction is therefore directional: positive coupling reflects structural architecture guiding function, while negative coupling reflects function operating in opposition to structural expectations, suggesting a complementary rather than symmetric relationship between the two states. Neutral coupling, which comprised the largest proportion of timepoints, suggests periods where functional dynamics operate independently of the specific structural priors used here, potentially reflecting flexible, polysynaptic signaling that transcends direct anatomical links (Zhao et al., 2021). The balance between these states and how they differ by sex offers a new window into the organization of the developing brain at this age.

We observed a robust dissociation in which males and females exhibit strong structure-function alignment (positive coupling). Females showed significantly stronger positive coupling in higher-order associative regions, specifically the inferior parietal lobule (IPL) and middle frontal gyrus (MFG). This finding aligns with established literature indicating that females often reach developmental peaks in cortical volume earlier than males (Giedd et al., 1997; Lenroot & Giedd, 2010) and possess proportionally larger volumes in language and associative cortices (Gur et al., 1999; Harasty et al., 1997). The stronger structural constraint in these regions is consistent with prior literature suggesting earlier integration of higher-order associative and executive control networks in females (Gong et al., 2011; Miller & Halpern, 2014). While prior work has linked these regions to social-cognitive development, our coupling-cognition findings support a more specific interpretation: higher positive coupling occupancy in these associative regions predicted better crystallized cognition, vocabulary and reading, in females but not males, suggesting that structural anchoring of these networks carries direct relevance for language-based cognitive performance at this age.

In contrast, males exhibited stronger positive coupling in sensorimotor and visuospatial regions (postcentral gyrus, precuneus). This pattern mirrors findings from adult structural connectome studies, which show that male brains are optimized for intra-hemispheric communication and sensorimotor coordination (Ingalhalikar et al., 2014), while female brains facilitate inter-hemispheric communication (Ingalhalikar et al., 2014). Furthermore, recent work on structure-function coupling in adults (Zhao et al., 2021) identified that males exhibit tighter coupling in "rich-club" connections, links between high-traffic structural hubs that tend to be concentrated in sensorimotor and subcortical regions, which form the densely interconnected core of the structural connectome. Females, by contrast, show stronger coupling in "feeder" connections, links between these central hubs and peripheral non-hub regions, which are predominantly located in higher-order associative cortices including the default mode, frontoparietal, and language networks. This alignment is therefore not coincidental: the sensorimotor anchoring we observe in males mirrors the rich-club architecture that is structurally dominant in males, while the associative anchoring in females mirrors the feeder architecture where female structural-functional coupling is strongest. Our results suggest these distinct organizational principles are already present and detectable in late childhood, extending prior adult findings into the developmental domain.

The cognitive moderation analysis delivers the proof-of-concept for dIMSC: if the coupling states which the method identifies are neurobiologically meaningful, they should predict cognition differently in males and females, in a manner consistent with known developmental patterns. This is precisely what we observe. Biological sex significantly modulated the relationship between dIMSC occupancy and cognitive performance, particularly in the positive coupling state. For females, higher occupancy in the positive state was associated with better performance in crystallized cognition (vocabulary, reading). These interactions were centered on the thalamus, cerebellum, and visual areas. This suggests that for females, accessing stored knowledge and verbal skills relies on a functional architecture that is tightly constrained by structural anatomy, particularly in thalamo-cerebellar and visual processing circuits. The thalamus, as a central relay hub connecting cortical and subcortical regions, may serve as a structural anchor for the integration of sensory and semantic information during crystallized cognition tasks, while the visual cortex involvement may reflect the visuospatial component of vocabulary and reading tasks. This is consistent with the “adaptive complementarity” hypothesis (Satterthwaite et al., 2015), where female advantages in verbal memory are supported by structural efficiencies across both thalamo-cortical relay circuits and temporo-parietal language networks. Together, these findings suggest that female crystallized cognition draws on structure-function coupling across multiple circuits and coupling states, rather than being localized to a single network.

For males, stronger positive coupling was associated with better performance in fluid cognition, specifically executive function and working memory. These interactions involved the hippocampus and paracentral lobule. This indicates that males who are better able to align their functional dynamics with their structural scaffold in these regions perform better on tasks requiring varying processing speed and inhibition. One possible explanation for why sensorimotor and visual regions specifically support fluid reasoning in males at this age relates to known sex differences in the neural architecture of visuomotor and spatial processing. Prior work has shown that males exhibit stronger intra-hemispheric connectivity in sensorimotor and visual networks during cognitively demanding tasks (Chen et al., 2024), and that this organizational pattern may confer a specific advantage for fluid reasoning tasks relying on visuospatial manipulation and pattern recognition, domains where male advantages are most consistently observed (Miller & Halpern, 2014). In females, the associative-anchored organization centered on temporo-parietal and frontal regions may provide a more efficient substrate for crystallized and verbal cognition, leaving the sensorimotor-visual circuit less specifically recruited for fluid reasoning at this developmental stage. These interpretations are necessarily speculative given the cross-sectional design, and longitudinal work will be needed to establish whether these coupling-cognition relationships reflect stable organizational features or transient developmental states. Interestingly, in the neutral state (weak coupling), males continued to show strong brain-behavior associations for working memory, whereas females did not. This may imply that males rely on a more “hard-wired” or structurally anchored strategy for executive functioning, even when functional dynamics are less constrained, whereas females may utilize more flexible, structurally independent functional states to achieve similar cognitive goals (Murray et al., 2021; Nini et al., 2017).

The stronger coupling in sensorimotor systems observed in males and in associative systems in females is consistent with theoretical models proposing that brain maturation proceeds along a sensorimotor-to-associative axis at different rates between sexes (Scheinost et al., 2015). Prior longitudinal work has linked prolonged synaptic pruning in lower-order networks to male development trajectories (Perrin et al., 2009), and earlier myelination in higher-order association cortices to female trajectories (De Bellis et al., 2001; Picci et al., 2023; Son et al., 2025). Our cross-sectional findings align with these patterns, though we cannot determine from this data whether these coupling differences reflect active maturational processes or stable sex-typical organizational features at this age. These distinct developmental trajectories underscore the importance of analyzing males and females separately in clinical studies.

While dIMSC offers a powerful new metric, this study is limited by its cross-sectional nature. Longitudinal analysis is required to determine if the male-sensorimotor/female-associative difference reflects a permanent sexual dimorphism or a transient developmental phase (i.e., males eventually “catching up” in associative coupling). Additionally, while we controlled for motion and volume, future work should explore how pubertal hormones (testosterone and estradiol) specifically drive these coupling shifts. Pubertal timing represents an important potential confound that was not directly addressed in the present study. Males and females in the 9–11 year age range differ not only in biological sex but also in pubertal stage, with females generally entering puberty earlier than males (Marshall & Tanner, 1969). Because pubertal hormones are known to regulate synaptic plasticity, myelination, and functional network reorganization in a sex-specific manner (Fung, Heinrichs-Graham, et al., 2022; Fung, Rahman, et al., 2022; Penhale et al., 2022; Sisk & Zehr, 2005), some observed coupling differences may reflect pubertal stage rather than sex per se. Regarding statistical power, while N=500 is sufficient to demonstrate proof-of-concept for the dIMSC framework, future work should replicate these coupling-cognition associations in larger samples to establish reproducibility, consistent with recommendations for brain-wide association studies (Marek et al., 2022).

## V. CONCLUSIONS

This study establishes dynamic inter-modality source coupling as a sensitive framework for mapping the time-varying interactions between brain structure and function. We provide evidence that the “rules” governing how function is constrained by structure differ fundamentally between sexes in childhood. Males exhibit a "sensorimotor-anchored" organization, where stronger structure-function coupling in motor and visual regions supports fluid reasoning. Females exhibit an “associative-anchored” organization, where coupling in parietal and frontal higher-order cortices supports crystallized cognition. Critically, the differential prediction of cognitive performance by dIMSC coupling states, varying systematically by sex in a manner consistent with known neurodevelopmental trajectories, serves as proof-of-concept validation for the method, demonstrating that the states dIMSC identifies are neurobiologically meaningful. These findings underscore the necessity of accounting for sex as a biological variable in neurodevelopmental research and establishes dIMSC as a sensitive framework for uncovering sex-linked patterns of dynamic brain organization with potential implications for future clinical and translational research.

## ABCD Data Acknowledgment

Data used in the preparation of this article were obtained from the Adolescent Brain Cognitive Development (ABCD) Study, held in the NIMH Data Archive (NDA). The ABCD Study is a multisite longitudinal study following more than 10,000 children recruited at ages 9–10 into early adulthood. The ABCD Study is supported by the National Institutes of Health and additional federal partners under award numbers U01DA041048, U01DA050989, U01DA051016, U01DA041022, U01DA051018, U01DA051037, U01DA050987, U01DA041174, U01DA041106, U01DA041117, U01DA041028, U01DA041134, U01DA050988, U01DA051039, U01DA041156, U01DA041025, U01DA041120, U01DA051038, U01DA041148, U01DA041093, U01DA041089, U24DA041123, and U24DA041147. ABCD Consortium investigators designed and implemented the study and/or provided data but did not necessarily participate in the analysis or preparation of this article. The views expressed in this article are those of the authors and do not necessarily represent the views of the NIH or ABCD Consortium investigators. The ABCD data repository grows and changes over time. The ABCD data used in this article came from ABCD Release 3.0, DOI: 10.15154/1519007.

