## Supplementary figures/tables for "Dynamic Inter-Modality Source Coupling Reveals Sex Differences in Children based on Brain Structural-Functional Network Connectivity: A Multimodal MRI Study of the ABCD Dataset"

**Supplementary Tables**

| **Term** | **Definition** |
| --- | --- |
| dIMSC | Dynamic inter-modality source coupling. Cross-correlation between the dFNC matrix and the SBM vector at each timepoint, quantifying the dynamic structure-function coupling strength. |
| Coupling state | Classification of a timepoint as positive, negative, or neutral based on the dIMSC correlation value relative to ρ=±0.1. |
| Positive coupling/state | Timepoints where dIMSC > 0.1; reflects strong alignment between functional networks activity and the underlying structural pattern. |
| Neutral coupling/state | Timepoints where -0.1 ≤ dIMSC ≤ 0.1; reflect periods where functional dynamics operate independently of the specific structural. |
| Negative coupling/state | Timepoints where dIMSC < -0.1; reflects periods where functional activity diverges from the structural pattern. |
| Occupancy rate | The proportion of total timepoints (out of 304) a given brain network spends in a particular coupling state, ranging from 0 to 1. Computed separately for each of the three coupling states (positive, neutral, negative) per subject per network. |

**Supplementary Table 1.** Glossary of key terminology used in the dIMSC pipeline. Definitions of core methodological terms introduced in the Dynamic Inter-Modality Source Coupling section of the Methods. All terms refer specifically to the dIMSC framework as implemented in this study.

| **Component** | **Positive state**  **(p-value)** | **Neutral state**  **(p-value)** | **Negative state (p-value)** |
| --- | --- | --- | --- |
| 1. Caudate | 0.130 | 0.0008*** | 0.149 |
| 2. Hypothalamus | 0.411 | 0.201 | 0.697 |
| 3. Putamen | 0.098 | 0.070 | 0.112 |
| 4. Caudate | 0.121 | 0.264 | 0.197 |
| 5. Thalamus | 0.169 | 0.172 | 0.545 |
| 6. Superior temporal gyrus | 0.097 | 0.002** | 0.012* |
| 7. Middle temporal gyrus | 0.265 | 0.064 | 0.301 |
| 8. Postcentral gyrus | 0.038* | 0.210 | 0.071 |
| 9. Left postcentral gyrus | 0.111 | 0.106 | 0.468 |
| 10. Paracentral lobule | 0.343 | 0.07 | 0.992 |
| 11. Right postcentral gyrus | 0.465 | 0.003** | 0.679 |
| 12. Superior parietal lobule | 0.711 | 0.700 | 0.314 |
| 13. Paracentral lobule | 0.559 | 0.088 | 0.015* |
| 14. Precentral gyrus | 0.995 | 0.008** | 0.325 |
| 15. Superior parietal lobule | 0.612 | 0.418 | 0.012* |
| 16. Postcentral gyrus | 0.093 | 0.270 | 0.057 |
| 17. Calcarine gyrus | 0.085 | 0.037* | 0.103 |
| 18. Middle occipital gyrus | 0.611 | 0.399 | 0.992 |
| 19. Middle temporal gyrus | 0.915 | 0.170 | 0.064 |
| 20. Cuneus | 0.996 | 0.121 | 0.128 |
| 21. Right middle occipital gyrus | 0.234 | 0.797 | 0.240 |
| 22. Fusiform gyrus | 0.064 | 0.099 | 0.404 |
| 23. Inferior occipital gyrus | 0.266 | 0.849 | 0.057 |
| 24. Lingual gyrus | 0.084 | 0.251 | 0.091 |
| 25. Middle temporal gyrus | 0.647 | 0.842 | 0.032* |
| 26. Inferior parietal lobule | 0.430 | 0.280 | 0.109 |
| 27. Insula | 0.101 | 0.243 | 0.086 |
| 28. Superior medial frontal gyrus | 0.265 | 0.0009*** | 0.104 |
| 29. Inferior frontal gyrus | 0.058 | 0.306 | 0.585 |
| 30. Right inferior frontal gyrus | 0.434 | 0.170 | 0.152 |
| 31. Middle frontal gyrus | 0.068 | 0.005** | 0.202 |
| 32. Inferior parietal lobule | 0.314 | 0.300 | 0.017* |
| 33. Left inferior parietal lobule | 0.026* | 0.111 | 0.488 |
| 34. Supplementary motor area | 0.684 | 0.551 | 0.430 |
| 35. Superior frontal gyrus | 0.054 | 0.522 | 0.316 |
| 36. Middle frontal gyrus | 0.040* | 0.678 | 0.106 |
| 37. Hippocampus | 0.143 | 0.062 | 0.500 |
| 38. Left inferior parietal lobule | 0.202 | 0.098 | 0.191 |
| 39. Middle cingulate cortex | 0.401 | 0.644 | 0.332 |
| 40. Inferior frontal gyrus | 0.088 | 0.071 | 0.097 |
| 41. Middle frontal gyrus | 0.058 | 0.084 | 0.283 |
| 42. Hippocampus | 0.316 | 0.646 | 0.057 |
| 43. Precuneus | 0.008** | 0.297 | 0.336 |
| 44. Precuneus | 0.391 | 0.032* | 0.061 |
| 45. Anterior cingulate cortex | 0.174 | 0.392 | 0.244 |
| 46. Posterior cingulate cortex | 0.491 | 0.086 | 0.293 |
| 47. Anterior cingulate cortex | 0.930 | 0.089 | 0.073 |
| 48. Precuneus | 0.868 | 0.099 | 0.090 |
| 49. Posterior cingulate cortex | 0.299 | 0.953 | 0.091 |
| 50. Cerebellum | 0.893 | 0.648 | 0.720 |
| 51. Cerebellum | 0.558 | 0.092 | 0.134 |
| 52. Cerebellum | 0.800 | 0.009** | 0.602 |
| 53. Cerebellum | 0.620 | 0.061 | 0.397 |

Supplementary Table 2. Sex differences in dIMSC occupancy rate across 53 brain components and three coupling states. P-values reflect independent samples t-tests comparing occupancy rate between females and males for each component across the positive, neutral, and negative coupling states. All p-values are FDR-corrected for multiple comparisons across 53 components and 3 coupling states. Significance levels: * 0.01 < p ≤ 0.05; ** 0.001 < p ≤ 0.01; *** p < 0.001. Components are organized according to the seven functional domains of the NeuroMark_fMRI_1.0 template: subcortical (1–5), auditory (6–7), sensorimotor (8–16), visual (17–25), cognitive control (26–33), default mode (34–49), and cerebellar (50–53).
